# Metabolic engineering of *Acinetobacter baylyi* ADP1 for naringenin production

**DOI:** 10.1101/2024.06.06.597799

**Authors:** Kesi Kurnia, Elena Efimova, Ville Santala, Suvi Santala

**Affiliations:** Faculty of Engineering and Natural Sciences, Tampere University, Hervanta Campus, 33720 Tampere, Finland

**Keywords:** Naringenin, *Acinetobacter baylyi* ADP1, coumaric acid, malonyl-CoA

## Abstract

Naringenin, a flavanone and a precursor for a variety of flavonoids, has potential applications in the health and pharmaceutical sectors. The biological production of naringenin using genetically engineered microbes is considered as a promising strategy. The naringenin synthesis pathway involving chalcone synthase (CHS) and chalcone isomerase (CHI) relies on the efficient supply of key substrates, malonyl-CoA and coumaroyl-CoA. In this research, we utilized a soil bacterium, *Acinetobacter baylyi* ADP1, which exhibits several characteristics that make it a suitable candidate for naringenin biosynthesis; the strain naturally tolerates and can uptake and metabolize coumarate, a primary compound in alkaline-pretreated lignin and a precursor for naringenin production. *A. baylyi* ADP1 also produces intracellular lipids, such as wax esters, thereby being able to provide an excess of malonyl-CoA for naringenin biosynthesis. Moreover, the genomic engineering of this strain is notably straightforward. In the course of the construction of a naringenin-producing strain, the coumarate catabolism was eliminated by a single gene knockout (Δ*hcaA*) and various combinations of plant-derived CHS and CHI were evaluated. The best performance was obtained by a novel combination of genes encoding for a CHS from *Hypericum androsaemum* and a CHI from *Medicago sativa,* that enabled the production of 18 mg/L naringenin in batch cultivations from coumarate. Furthermore, the implementation of a fed-batch system led to a significant 3.7-fold increase (66 mg/L) in naringenin production. These findings underscore the potential of *A. baylyi* ADP1 as a host for naringenin biosynthesis as well as advancement of lignin-based bioproduction.

## 1. Introduction

Flavonoids are an important class of natural products widely found in the plant kingdom (Panche et al., 2016; Pandey et al., 2016). They belong to a class of plant polyphenols, comprising a family of more than 9000 compounds such as flavones, flavanols, isoflavones, anthocyanins, and chalcones (Panche et al., 2016; Zhang et al., 2021; Li et al., 2022). They possess a wide array of valuable applications in the realms of health and pharmaceuticals, encompassing attributes such as anticancer, antioxidant, anti-inflammatory, and antiviral properties, along with recognized neuroprotective and cardioprotective effects (Salehi et al., 2019; Zhang et al., 2021; Cai et al., 2023). Among various classes, one of the important flavonoids is a flavanone naringenin which occupies a central position as the primary C_15_ intermediate in the flavonoid biosynthesis pathway (Hwang et al., 2021; Zhang et al., 2021). This compound has the carbon framework common to flavonoids (C6−C3−C6) and functions as a precursor to other flavonoids, including flavanones, flavanols, and isoflavonoids (Li et al., 2022; Liu et al., 2022). In addition, a recent in vitro test study has indicated that naringenin could potentially offer a promising strategy for the treatment of COVID-19 (Tutunchi et al., 2020; Clementi et al., 2021).

Traditionally, the production of flavonoid and flavonoid-derived compounds has relied heavily on their isolation from plants, necessitating extensive separation and purification efforts (Koopman et al., 2012a; Ye et al., 2023). This approach has proven to be neither cost-effective nor sustainable as a source of flavonoids. Moreover, the conventional method has faced limitations due to low yield obtained from natural sources (Karim et al., 2018; Hwang et al., 2021). To overcome these challenges, numerous attempts have been undertaken to synthesize naringenin in microbial systems.

Due to the abundancy of genetic tools, conventional workhorses such as *Escherichia coli*, *Corynebacterium glutamicum,* and *Saccharomyces cerevisiae*, have been previously employed in the production of flavonoids (Santos et al., 2011; Koopman et al., 2012; Wu et al., 2014; Milke et al., 2019; Wu et al., 2022). In microbial hosts, naringenin can be produced from one molecule of *p*-coumaroyl-CoA and three molecules of malonyl-CoA via a naringenin chalcone intermediate (Gao et al., 2020; Liu et al., 2022). For instance, Zhou et al. (2021) produced naringenin from glucose and glycerol in *E. coli* by introducing a chalcone synthase (CHS) from *Petunia hybrida* and a chalcone isomerase (CHI) from *Medicago sativa.* Cells were also supplemented with palmitate and stearate to increase the amount of malonyl-CoA obtained from their degradation. In another study by Milke et al.(2019), CHS and CHI from *P. hybrida* were expressed in *C. glutamicum* to produce naringenin from glucose. To increase the malonyl-CoA availability, they regulated the fatty acid synthesis and reduced the flux into the tricarboxylic acid cycle (TCA cycle) by metabolic engineering.

Due to the challenges related to the malonyl-CoA availability, the potential of oleaginous microorganisms in the production of flavonoids is increasingly investigated. These organisms can produce lipids exceeding 20% of their cell dry weight, which implies the potential to overproduce malonyl-CoA. Recently, Zan et al., (2024) engineered a non-model organism *Mucor circinelloides* to produce naringenin by a heterologous pathway. Despite the obtained titer was relatively low, 2.2 mg/L, the work nicely demonstrates the potential of leveraging the naturally active anabolic pathways for malonyl-CoA synthesis.

Many of the previously described naringenin production systems rely on the utilization of glucose and/or glycerol as substrates for the synthesis (Santos et al., 2011; Koopman et al., 2012; Hwang et al., 2023). However, the key naringenin precursor, coumarate, represents a major compound of alkaline-pretreated lignin (APL), which could serve as a renewable feedstock for biocatalysts (Vardon et al., 2015; Incha et al., 2020; Weiland et al., 2022). Despite being the second most abundant biopolymer from terrestrial plants, lignin remains significantly underutilized (Becker and Wittmann, 2019; Yu et al., 2023). For instance, chemical pulping operations generate more than 50 million tons of technical lignin as by-products, with most of this being burnt as low-quality fuels (less than $50/dry ton) (Balakshin et al., 2021). Consequently, utilizing lignin-derived aromatics for microbial naringenin production could serve as a sustainable means to utilize natural resources while concurrently enhancing the competitiveness of the lignocellulose-based biorefinery processes.

*Acinetobacter baylyi* ADP1 emerges as a promising candidate for the biological conversion of lignin-derived compounds, due to its capability to uptake and utilize aromatic substances through the *β*-ketoadipate pathway (Harwood and Parales, 1996; Bleichrodt et al., 2010; Salvachúa et al., 2015; Luo et al., 2019; Biggs et al., 2020). The strain can also tolerate and grow on significantly high concentrations of lignin-related aromatics (Luo et al., 2022b). Moreover, we have previously demonstrated *A. baylyi* ADP1 to be a potential host for the overproduction of intracellular storage lipids, namely wax esters (Santala et al., 2018; Salmela et al., 2019; Luo et al., 2020; Santala et al., 2021). This pathway can be associated with the potentially high supply of malonyl-CoA for syntheses pathways. Additionally, *A. baylyi* ADP1 is highly engineerable and naturally competent, which facilitates swift and targeted genomic manipulations (Elliott and Neidle, 2011; Biggs et al., 2020; Santala and Santala, 2021) and evolution-based engineering (Tumen-Velasquez et al., 2018; Bedore et al., 2023). Taken together, these features highlight ADP1 as an ideal host for the production of flavonoid-like products from sustainable resources.

In this work, we established naringenin production in *A. baylyi* ADP1 (Figure 1). We exploited the strain’s native ability to utilize coumarate as a substrate and established a non-native pathway for naringenin synthesis. In addition, we implemented a fed-batch process to improve naringenin production, resulting in a notable improvement in naringenin titer. This study demonstrates *A. baylyi* ADP1 as a promising host for biological valorization of lignin-related compounds into high-value flavonoids.

**Fig 1.**
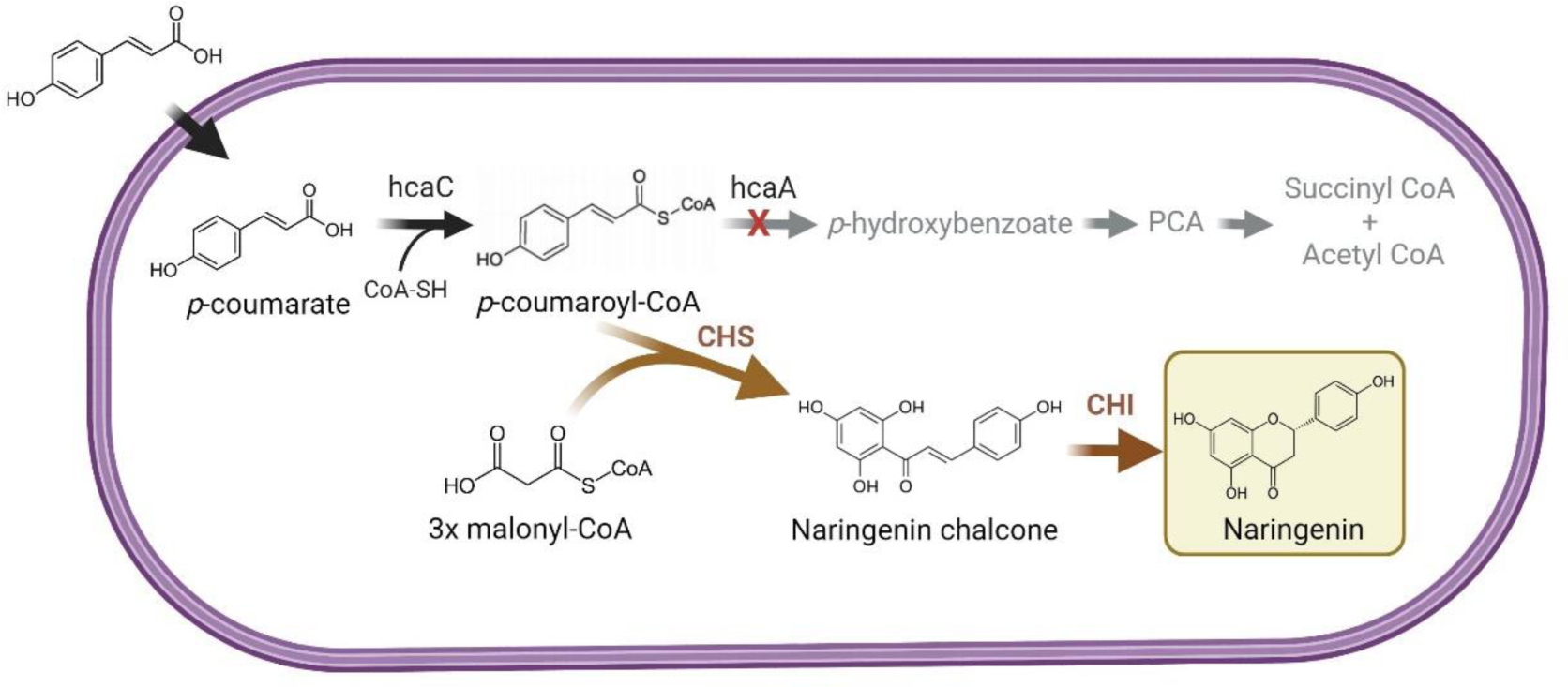
Schematic representation of heterologous production of naringenin in *A. baylyi* ADP1. Black arrows indicate the native pathways. Grey arrows indicate pathway abolished by deletion of a bifunctional hydratase/lyase (*hcaA*), and brown arrows indicate naringenin production pathway. Abbreviations: protocatechuate (PCA), chalcone synthase (CHS), chalcone isomerase (CHI).

## 2. Materials and methods

### 2.1 Bacterial strains and culture conditions

*Escherichia coli* XL1-Blue (Stratagene, USA) was used for the plasmid constructions and amplifications. Strains were maintained on Lysogeny broth (LB) or LB agar containing antibiotics for plasmid selection. *A. baylyi* ADP1 (DSM 24193, DSMZ, Germany) was used for naringenin production and pathway engineering. *A. baylyi* ADP1 and derived strains were routinely grown aerobically at 30 °C in LB or Minimal salts medium (MSM). For grown-on plates, 1.5% agar (w/v) was added. Corresponding antibiotics were added to the media when necessary (100 μg/mL ampicillin, 25 μg/mL chloramphenicol, 50 μg/mL kanamycin, and gentamycin 15 μg/mL). All growth and production experiments were performed using MSM (per Liter; 3.88 g K_2_HPO_4_, 1.63 g NaH_2_PO_4_, 2.0 g (NH_4_)_2_SO_4_, pH 7.0). The MSM media was supplemented with a trace elements solution (per Liter; 10 mg L^−1^ ethylenediaminetetraacetic acid (EDTA), 0.1 g/L MgCl_2_.6H_2_O 2 mg/L ZnSO_4_7H_2_O, 1 mg/LCaCl_2_·2H_2_O, 5 mg/LFeSO_4_·7H_2_O, 0.2 mg/L, Na_2_MoO_4_·2H_2_O, 0.2 mg/L CuSO_4_·5H_2_O, 0.4 mg/L CoCl_2_·6H_2_O, 1 mg/L MnCl_2_·2H_2_O). The stock solution of coumarate was prepared with a concentration of 200 mM (pH 8.2-8.3) (Luo et al., 2022b). Cerulenin (Sigma Aldrich) was dissolved in ethanol as previously described (Schwanemann et al., 2023). Sodium malonate and naringenin were purchased from Sigma Aldrich.

### 2.2 Genetic modifications

Primers were synthesized by Thermo Scientific and routine PCR amplifications were carried out using Phusion High Fidelity DNA Polymerase (New England Biolabs, UK). Chromosomal modifications were engineered in *A. baylyi* ADP1 by natural transformation of recipient strains with linear DNA fragments as described previously (Luo et al., 2020, 2022a). First, regions of approximately 500 bp-1000 bp upstream (R1) and downstream (R2) of the target genes were amplified from the genomic DNA of *A. baylyi* ADP1. The R1 and R2 were constructed with tdk-kanR cassette by OE-PCR and transformed to *A. baylyi* ADP1 as described previously (De Berardinis et al., 2008). The rescue cassette (1-2 kb) was introduced to generate a clean knock-out. *A* successful deletion in *A. baylyi* ADP1 was screened on negative selection LB plates with AZT + 0.1% glucose. Genotypes were then confirmed by colony PCR with OneTaq (New England Biolabs, UK) and selected for antibiotic sensitivity. Detailed information regarding all strains and primers used in this study is provided in Table S1 and Table S2.

### 2.3 Pathway construction

Genes encoding malonyl-CoA synthetase from *Rhizobium trifolii* (matB, KF765783.1), chalcone synthase (CHS) from *Petunia X hybrida* (PhCHS, KF765781.1), *Huperzia serrata* (HsPKS1, DQ979827.1), *Hypericum androsaemum* (HaCHS, AF315345.1) and genes encoding chalcone isomerase (CHI) from *Pueraria lobata* (PlCHI, D63577.1) and *Medicago sativa* (MsCHI, KF765782.1) were used in this study. Those genes were synthesized and codon-optimized for ADP1 by GenScript Biotech (Netherlands) with appropriate restriction sites and ribosomal binding sites. The naringenin plasmids were constructed as follows. First, each CHS gene was cloned to pBAV1C-chn (Luo et al., 2019) with BioBrick assembly standard using restriction sites XbaI/SpeI and PstI. Subsequently, the resulting plasmid was digested with XbaI and PstI. The CHI gene was cloned downstream of the CHS gene, resulting in the creation of plasmids pNAR1, pNAR2, pNAR3, pNAR4, pNAR5, and pNAR6.

The integration of *matB* from *Rhizobium trifolii* into the genome was facilitated by previously described gene integration cassette (Santala et al., 2011a; Lehtinen et al., 2018). The *matB* gene was cloned to integration cassette by utilizing NdeI/XhoI restriction sites. This cassette contains two antibiotic markers, Cm^R^ and Kan^R^, allowing selection in *E. coli* and *A. baylyi* ADP1, respectively. Furthermore, this cassette enables the replacement of the genes *poxB* (ACIAD3381), metY (ACIAD3382) and *acr1* (ACIAD3383) with the *matB* gene. Notably, the genes *poxB* and *metY* were targeted for deletion due to their recognized neutrality for deletion, as prior study have shown their removal does not adversely impact the growth of ADP1 (Santala et al., 2011b; Luo et al., 2020). The constructed plasmids were transformed to *E. coli* XL-1 cells and transformants were selected on LB plates with 25 µg/ml chloramphenicol. Verified plasmids were then transformed into *A. baylyi* ADP1 using natural transformation. All plasmid inserts and integration were verified by sequencing (Macrogen, the Netherlands). The gene sequences can be found in Supplementary Table S4.

### 2.4 Small-scale naringenin production and fed-batch fermentation

To monitor naringenin production, single colonies were inoculated into 14 mL tubes with 5 mL MSM containing 50 mM gluconate and 0.2% casamino acids. The precultures were incubated at 30 ^◦^C at 300 rpm. After overnight cultivation, the precultures were diluted and transferred to a 100 mL flask containing 20 mL MSM with 50 mM gluconate and 0.2% casamino acids at an initial OD_600_ of 0.7-0.8. Coumarate was added at a concentration 2.5 mM, and cyclohexanone at 5 μM was used to induce heterologous pathway expression. All cultivations were performed in triplicate and were allowed to proceed for 72 hours. ADP1 containing empty plasmid was used as a control.

The fed-batch culture was performed in a 250 mL mini bioreactor (Applikon Biotechnology, The Netherlands) with a 50 mL initial volume. Prior to inoculation, the process parameters were set at a 100% dissolved oxygen, 200 rpm stirrer speed, and temperature 30 °C. A fresh colony of the constructed derivatives was grown into MSM containing 50 mM gluconate and 0.2% casamino acids. Subsequently, the culture was transferred to a bioreactor at an initial OD_600_ 0.7-0.8. The feed solution (50 mM gluconate, 0.2% casamino acids, and 3.2 mM coumarate in MSM) was fed at 3.7 mL per hour for 48h. The cyclohexanone was added at a concentration 0.5 μM to induce the pathway expression. For the experiment with pH control, the pH of the fermentation was maintained at 7.5 through the addition of 1 M H_3_PO_4_. Additionally, if necessary, the 10% antifoam A (Fluka Analytical) was added. Samples (2 mL) were taken in aseptic conditions throughout the duration of the cultivation to measure cell growth, coumarate, and naringenin. The bioreactor experiments were conducted in duplicate.

### 2.5 Analytical methods

Optical density was measured at 600 nm using an Ultrospec 500 Pro (Amersham Biosciences). For measurement of naringenin, the fermentation broth was mixed with an equal volume of ethyl acetate, vortexed for 30 s, and rotated at room temperature for 2h. After centrifugation, the upper layer was collected, and the solvent was allowed to evaporate overnight. The dried samples were then resuspended in methanol for analysis with HPLC. For measurement of coumarate, the culture was centrifuged at 20,000g for 5 min. The supernatant was taken and filtered with syringe filters (CHROMAFIL® PET, PET-45/25, Macherey–Nagel, Germany). Identification of naringenin and coumarate was performed using liquid chromatography coupled to photodiode-array detection (Shimadzu, USA). Luna C18(2) 100Å 4.6 x 150 mm was used as the column and placed at 40 °C. An isocratic elution program was applied at a flow rate 1 mL/min, using 60% (0.1% formic acid) and 40% (MeOH). The sample injection was set to 5 μL and the signal was detected at 288 nm at the retention time of approximately 4.29 min (coumarate) and 16.44 min (naringenin). The concentrations were determined through the use of the corresponding chemical standards (Sigma).

## 3. Result and Discussion

### 3.1 *Acinetobacter baylyi* ADP1 as a naringenin production platform

*A. baylyi* ADP1 possesses multifaceted aromatic catabolic capability, notably displaying efficient utilization of coumarate as a precursor for naringenin (Bleichrodt et al., 2010; Harwood and Parales, 1996). In addition, the ability to accumulate intracellular lipids associated with high supply of malonyl-CoA renders *A. baylyi* ADP1 as an optimal candidate for the production of flavonoids, such as naringenin. As naringenin has reported to have antimicrobial effects (Lather et al., 2020; Cai et al., 2023), we first wanted to assess the tolerance of *A. baylyi* ADP1 towards naringenin by culturing the strain in a mineral salt medium (MSM) with 50 mM gluconate and 0.2% casamino acids with varying concentrations of naringenin (Fig. 2A). The cells were able to grow in the presence of all studied naringenin concentrations; growth was not affected by up to 50 mg/L of naringenin, while growth was somewhat inhibited in the presence of 200 mg/L of naringenin. We also confirmed that *A. baylyi* ADP1 is not able to degrade naringenin (Fig. 2B); no obvious decrease in naringenin concentration was observed after 48h fermentation. The above results indicated it is possible to harness *A. baylyi* ADP1 for naringenin production.

**Fig. 2.**
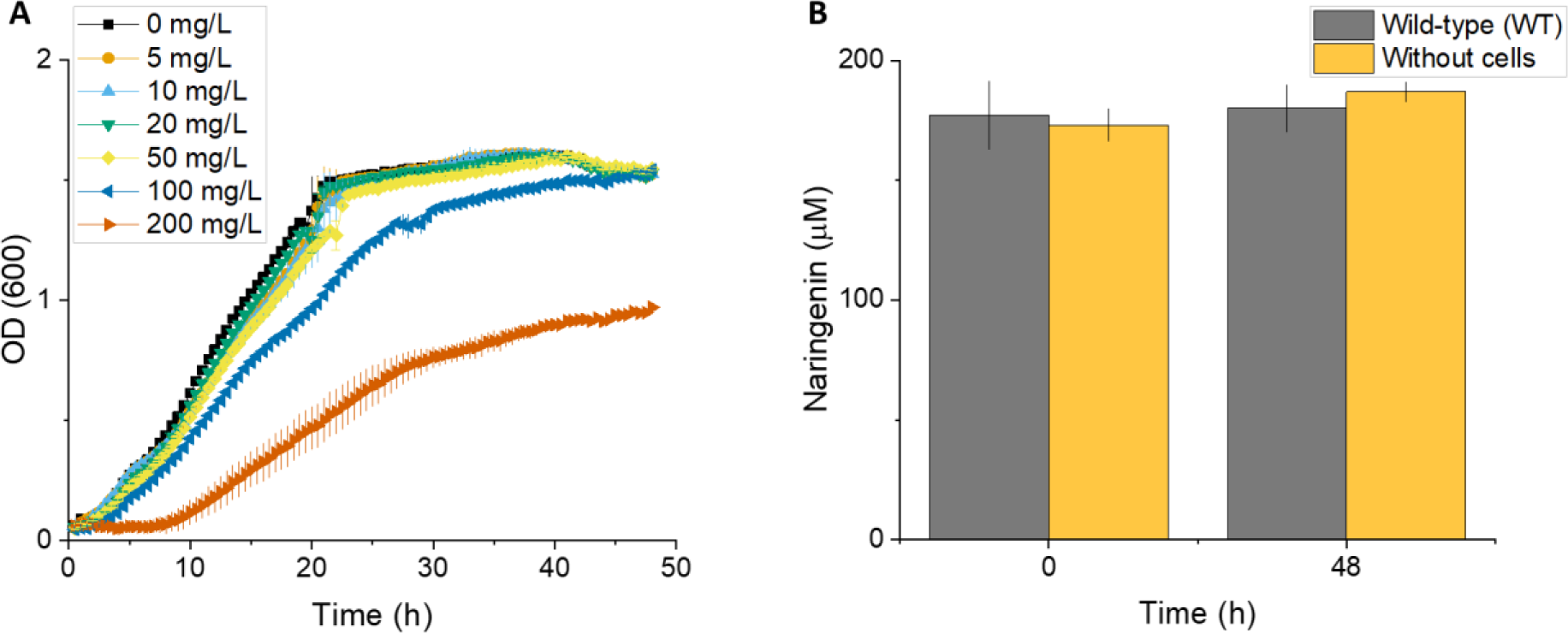
Assessment of *A. baylyi* ADP1 growth in the presence of naringenin. A) Growth of *A. baylyi* ADP1 wild-type (WT) in the presence of an increasing concentration of naringenin in MSM with 50 mM gluconate and 0.2% casamino acids. B) Stability of naringenin in *A. baylyi* ADP1 WT culture. Cells were incubated in MSM with 200 uM (54.45 mg/L) naringenin. Error bars represent the mean ± s.d of three biological replicates.

### 3.2 Screening an optimal naringenin production module

Naringenin is the starting point for many flavonoid functionalization chemistries (Hwang et al., 2021; Zhang et al., 2021). It is synthesized via the phenylpropanoid metabolic pathway in plants, requiring one molecule of *p*-coumaroyl-CoA and three molecules of malonyl-CoA, catalyzed by the enzyme chalcone synthase. Our aim was to construct the naringenin production pathway from coumarate via chalcone intermediate. First, given that *A. baylyi* ADP1 naturally generates coumaroyl-CoA as part of its coumarate catabolism, we performed a genetic knockout of the downstream gene in the native catabolic pathway. Specifically, we deleted the *hcaA* gene, which encodes a bifunctional hydratase/lyase responsible for converting the thioester derivative to an aldehyde intermediate (Parke and Ornston, 2004). This genetic manipulation was executed with the intention of preventing *A. baylyi* ADP1’s consumption of these necessary precursors. Subsequent cultivation experiments confirmed that the *A. baylyi* ADP1 Δ*hcaA,* designated as ASA800 was incapable of utilizing coumarate as the sole carbon source (Fig. S1).

Subsequently, we proceeded to establish a synthetic pathway and confirmed the viability of employing ASA800 as the host organism for flavonoid production. To construct the naringenin production module, two genes, chalcone synthase (CHS) and chalcone isomerase (CHI) are required. In this study, we introduced CHS originating from various plants, *Petunia X hybrida* (PhCHS)*, Huperzia serrata* (*Hs*PKS1), and *Hypericum androsaemum* (*Ha*CHS), that accept *p*-coumaroyl-CoA as a substrate (Liu et al., 2003; Wu et al., 2014; Palmer et al., 2020). In addition, CHI which is responsible for the further isomerization of chalcone to flavanone, was selected from *Petunia lobata* (PlCHI) and *Medicago sativa* (MsCHI). The CHS-CHI genes were previously expressed for naringenin or other flavonoids production in *E. coli* (Wu et al., 2014; Van Brempt et al., 2022), *Saccharomyces cerevisiae* (Lyu et al., 2019), *Yarrowia lipolytica* (Palmer et al., 2020) and *Corynebacterium glutamicum* (Kallscheuer et al., 2016). Here the assembly of these orthologues resulted in the generation of six recombinant pathways in which HaCHS – MsCHI represents a novel combination (Fig. 3A). To ensure robust expression of the heterologous genes, genes were expressed in the plasmid pBAV1C-chn under cyclohexanone-inducible promoter ChnR/P_ChnB_ (Luo et al., 2019). The constructed plasmids (pNAR1-pNAR6) were then transformed to ASA800, resulting six naringenin-producing strains, designated as ASA801-ASA806, respectively. The ADP1 Δ*hcaA* harboring empty plasmid pBAV1C-chn (ASA807) was used as a control. The ASA801-ASA807 were then cultivated in MSM with 50 mM gluconate, 0.2% casamino acids, and 2.5 mM coumarate for 72 h. All tested combinations were found to be functional and produce naringenin. However, the different combinations of naringenin production modules resulted in varying naringenin titers ranging from 6 mg/L to 18 mg/L (Fig. 3B). The highest titers were reached already after 24 h cultivations by cells harboring pNAR3 (ASA803) and pNAR6 (ASA806), yielding 17 mg/L and 18 mg/L, respectively. Interestingly, both these production strains contained HaCHS, indicating that CHS mainly dictates the efficiency of the naringenin production module in the studied combinations. Thus, these results demonstrated the feasibility of using *A. baylyi* ADP1 as the chassis for de novo synthesis of naringenin.

**Fig. 3.**
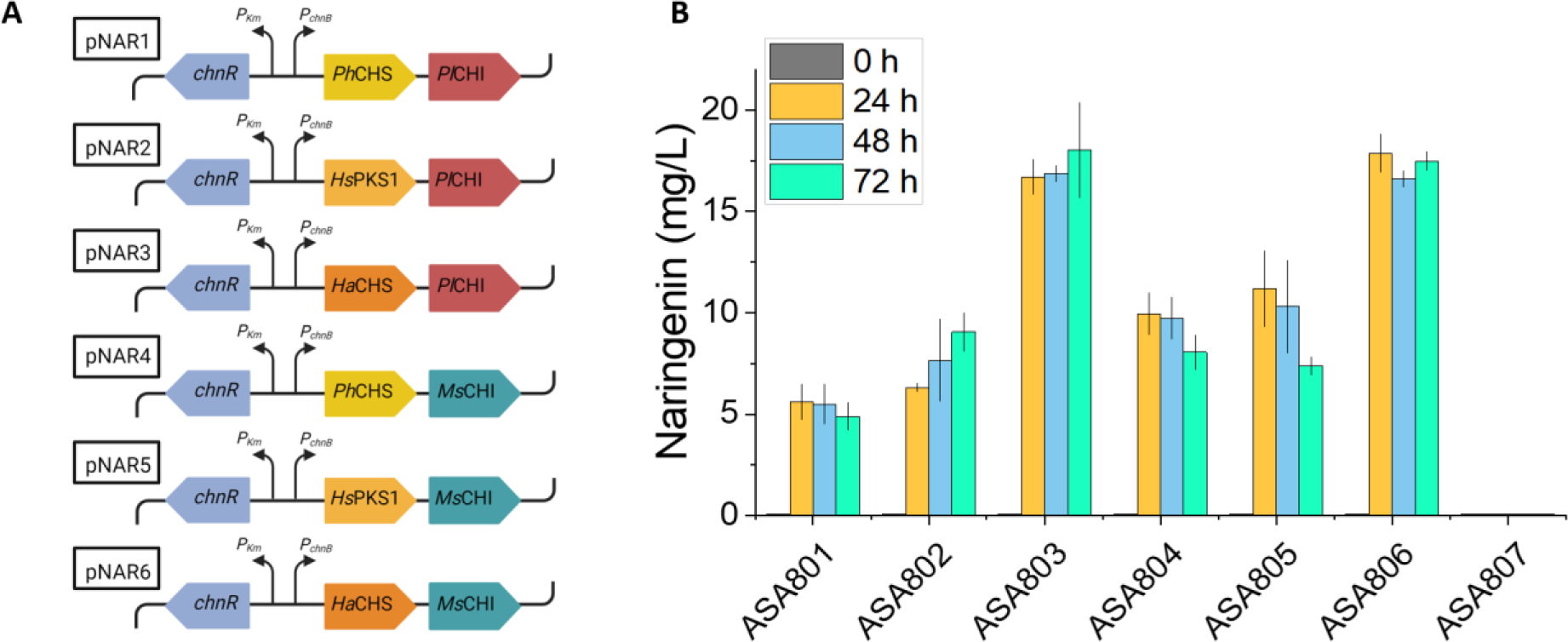
Naringenin production modules in constructed strains. A) Plasmid construction for CHS and CHI expression from different plant species. B) Naringenin production by ASA801-ASA807 in flasks supplemented with 2.5 mM coumarate. pBAV1C-chn was used as an empty plasmid. The expression was regulated by inducible promoter P*_ChnB_*. Error bars represent the mean ± s.d of three biological replicates.

### 3.3 Toxicity effect caused by coumarate in ADP1 naringenin producing strain

In previous research (Parke and Ornston 2004), it was highlighted that the deletion of the hydratase/lyase gene *hcaA*, abolishes the ability of *Acinetobacter* cells to grow on the three hydroxycinnamates (caffeate, *p*-coumarate, and ferulate). Furthermore, this genetic alteration also inhibits cell growth in the presence of these compounds even in concentrations as low as 100 μM (caffeate), 10 μM (*p*-coumarate) and 1 mM (ferulate) (Parke and Ornston, 2003, 2004). Similar toxicity effects by coumaroyl-CoA have been observed also in *Pseudomonas putida* KT2440 (Incha et al., 2020) and *Saccharomyces cerevisiae* (Liu et al., 2022). Thus, we investigated the impact of coumarate concentration on our naringenin production strains ASA803 and ASA806. The strains were tested with two different coumarate concentration (2.5 mM and 10 mM) and naringenin levels were assessed after 24 hours of cultivation. We observed that coumarate concentration of 10 mM resulted in decreased naringenin production in both ASA803 and ASA806, indicating a toxic effect of this compound (Fig. 4A). This observation supports the notion that a high concentration of the coumaroyl-CoA intermediate is toxic to ASA800.

**Fig 4.**
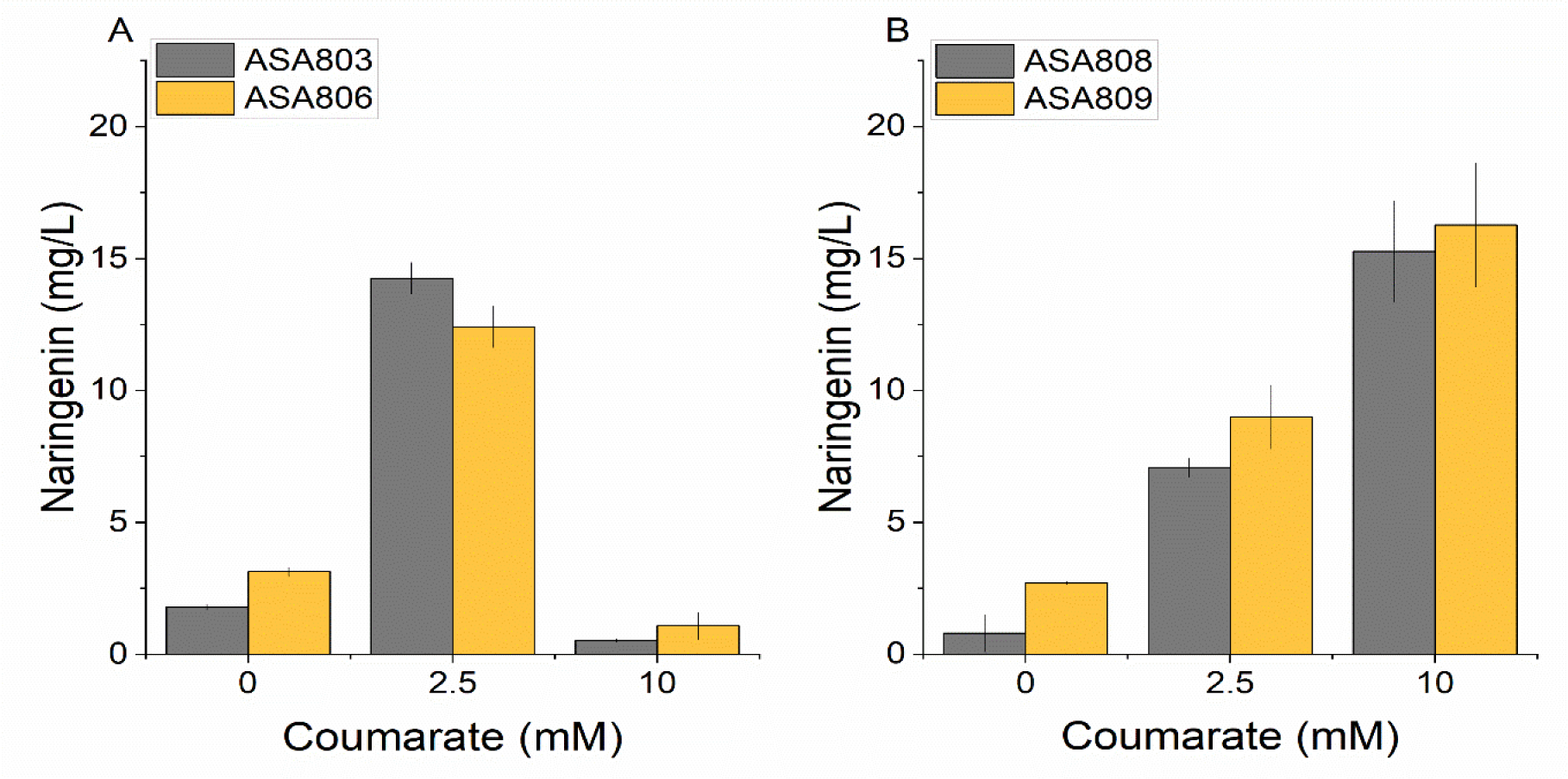
Effect of coumarate concentration on naringenin production in A) ASA803 and ASA806, B) ASA808 and ASA809. Cells were cultivated in MSM with 50 mM gluconate, 0.2% casamino acids, and different concentration of coumarate. Gene expressions were induced with 5 μM cyclohexanone. Samples for analysis were taken after 24 h. Error bars represent the mean ± s.d of three biological replicates.

To further investigate the potentially negative effect of the *hcaA* deletion, we conducted a test with the wild-type (WT) ADP1 carrying pNAR3 (ASA808) and pNAR6 (ASA809) for comparison (Fig. 4B). We hypothesized that a potential ‘pulling effect’ generated by native coumarate utilization pathway might have positive effect on the naringenin accumulation. Indeed, we observed that with high coumarate concentration (10 mM), a higher naringenin titer was obtained compared with the Δ*hcaA* strains. However, with lower coumarate concentration (2.5 mM) the titer was lower with the WT strains, indicating that a significant amount of coumarate is channeled to the catabolic pathway instead of the naringenin synthesis pathway. Thus, for further experiments, we chose to continue with ASA803 and ASA806 using lower concentrations of coumarate (2.5 mM), for which the tolerance was found to be sufficient.

### 3.4 Effect of deletion of malonate utilization pathway and addition of FAS cerulenin on naringenin production

Malonyl-CoA constitutes a vital precursor for the biosynthesis of flavonoids (Milke et al., 2019; Wu et al., 2015). It serves as a co-substrate for CHS, and its substantial presence plays crucial role in naringenin production (Wu et al., 2022; Xu et al., 2011). Therefore, the availability of malonyl-CoA can be the principal limiting factor in the flavonoid synthesis. Several efforts have been made to increase the pool of this important precursor molecules. For example, Wu et al., 2022 introduced *matB* and *matC* genes from *Rhizobium trifolii* into *Corynebacterium glutamicum*, resulting in a remarkable 35-fold increase in naringenin production. In this context, we investigated two strategies for increasing the supply of malonyl-CoA to improve naringenin yields. In the first approach, we obstructed the malonate degradation pathway in ADP1 and introduced *matB* gene from *Rhizobium trifolii*, the enzyme responsible for the conversion of malonate to malonyl-CoA. In *A. baylyi* ADP1, a transcriptional regulation mechanism, characterized by a cluster of *mdc* genes, facilitates the utilization of malonate as a carbon source (Stoudenmire et al., 2017). Thus, in order to prevent further malonate degradation, the malonate pathway in ADP1 was eliminated, involving the deletion of *mdcA, mdcB, mdcC, mdcD, mdcE, mdcG*, and *mdcH*. Additionally, we incorporated *matB* from *R. trifolii* into the genome using the previously described gene integration cassette (Lehtinen et al., 2018). This cassette also removes the strain’s ability to produce wax ester due to the deletion of the gene encoding for the fatty acyl-CoA reductase *acr1* (ACIAD3383) essential for wax ester synthesis (Santala et al., 2014). Subsequently, we conducted experiment to assess naringenin production in MSM medium supplemented with 50 mM gluconate, 0.2% casamino acid, 2.5 mM coumarate and 15 mM sodium malonate. Regrettably, the Δ*hcaA* Δ*mdcABCDEGH* Δ*ACIAD3383-3381::matB* harboring pNAR3 (ASA810) failed to grow on this medium at the studied conditions. We further investigated what caused the growth defect by integrating the *matB* into ADP1 Δ*mdcABCDEGH* and tested in MSM medium supplemented with 50 mM gluconate, 0.2% casamino acid, 2.5 mM coumarate and 15 mM sodium malonate. The result showed a minor growth defect in Δ*mdcABCDEGH* Δ*ACIAD3383-3381::matB* pNAR3 (ASA811) compared to ASA808 (Fig. S2). Furthermore, integration *matB* did not increase naringenin production. It appears that these genetic modifications may affect the primary metabolism of ADP1 and would thus require more investigation. Consequently, the outcome suggests that eliminating the malonate degradation pathway might not constitute an efficient strategy for improving naringenin production in ADP1.

The second approach involved the utilization of cerulenin, a compound known for its inhibitory action on fatty acid biosynthesis. Cerulenin binds to the b-ketoacyl-acyl carrier protein (ACP) synthase of the fatty acid machinery, thereby restricting the depletion of malonyl-CoA for fatty acid biosynthesis (Santos et al., 2011). Supplementation of cerulenin has proven improved resveratrol in *C. glutamicum* (Kallscheuer et al., 2016)*, P. taiwanensis* (Schwanemann et al., 2023), and naringenin in E. coli (Santos et al., 2011). Given the identical CHS gene in both pNAR3 and pNAR6 and the high cost associated with cerulenin, in this study, we conducted experiment with only ASA803. As depicted in Fig. S3A, the growth of our strain remained unaffected by a 50 μM concentration of cerulenin, a factor advantageous for both naringenin yield and productivity. In prior research, it was observed that cerulenin supplementation negatively impacted the growth in *C. glutamicum* (Kallscheuer et al., 2016) and *E. coli* (Leonard et al., 2008) while producing higher amount of flavonoids. However, ASA803 exhibited lower naringenin production in the presence of cerulenin (Fig. S3B). This parallels the finding documented by Dunstan et al., (2020), regarding the CHS from *Camelia sinensis* (Tea), showing similar outcome (Dunstan et al., 2020). Similar findings were also reported in *Streptomyces albidoflavus* J1074, where cerulenin failed to increase naringenin production (Ye et al., 2023). Ferrer et al., 1999 reported that cerulenin is a potent irreversible inhibitor of CHS through its binding to Cys-164, thereby restricting access to the coumaroyl-binding pocket (Ferrer et al., 1999). Hence, it is plausible that instead of significantly affecting fatty acid biosynthesis, cerulenin might bind to CHS, impending naringenin biosynthesis in *A. baylyi* ADP1.

### 3.5 Naringenin production in a bioreactor

CHS and CHI activity are recognized to exhibit pH-dependent behavior (Jez and Noel, 2002; Dunstan et al., 2020). Hence, for the purpose of further characterizing the naringenin-producing strain under controlled conditions, these strains were cultured in a 250 mL bioreactor set at a pH 7.5. As a comparative measure, a cultivation in a bioreactor without pH controlled was also subjected to testing. Further, to avoid the adverse impact of high concentration of coumarate and obtain a higher titer of naringenin, a fed-batch approach was implemented. Additionally, in contrast to the flask experiment, a lower inducer concentration (0.5 µM instead of 5 µM of cyclohexanone) was employed (Fig. S4). As shown in Fig. 5A and 5B, under pH-controlled conditions, the ASA803 and ASA806 cultures yielded 37 mg/L and 34 mg/L of naringenin, respectively. This represented a significant increase in naringenin production compared to the flask experiment, with a 2.2-fold and 1.9-fold enhancement. Surprisingly, when pH was not controlled, higher naringenin titer was observed in ASA803 and ASA806, producing 48 mg/L and 60 mg/L, respectively (Fig. 5C and Fig. 5D). Previous studies demonstrated that naringenin chalcone could be converted to naringenin at higher pH yielding higher naringenin production (Hwang et al., 2003; Wu et al., 2014). Furthermore, our result shows that MsCHI from *M. sativa* resulted in an increase naringenin production compared to PhCHI from *P. hybrida*. This finding corroborates the previous results where it was observed that the replacement of PhCHI with MsCHI from *M. sativa* led to an enhanced production of pinocembrin, naringenin, and eridictyol (Leonard et al., 2007). Furthermore, to the best of our knowledge, pNAR6 (HaCHS – MsCHI) represents a novel combination as this specific configuration has not been previously expressed in other microbial hosts.

**Fig. 5.**
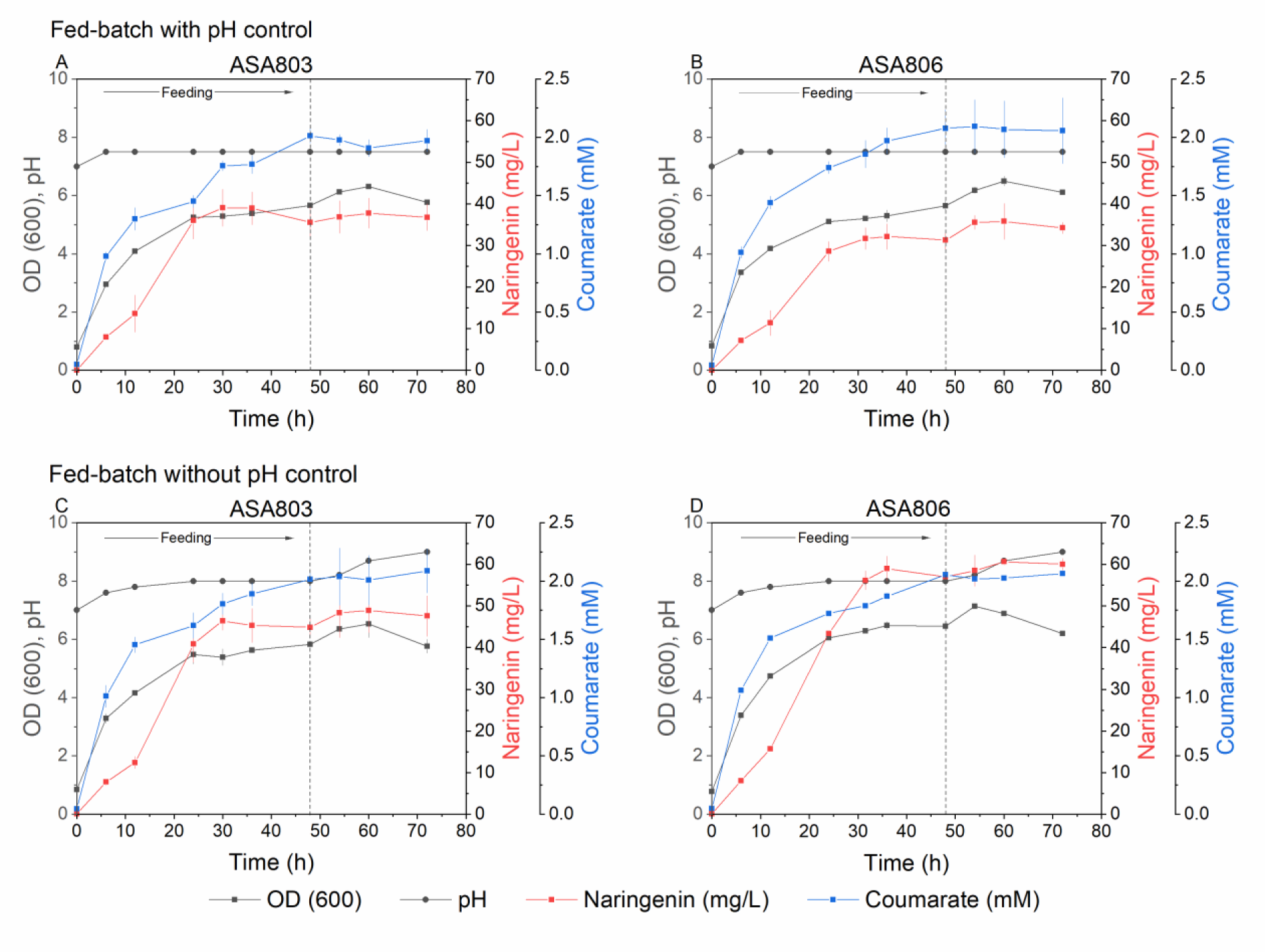
Naringenin production in ASA803 and ASA806 in fed-batch cultivation with pH control (A) and without pH control (B). Cultures initially contained 50 µM of coumarate, 50 mM gluconate, and 0.2% casamino acids. Cultures were fed with a medium containing 50 mM gluconate, 0.2% casamino acids, and 3.2 mM coumarate for 48h (feeding rate 3.7 mL/h). Error bars represent the mean ± s.d of two biological replicates.

Next, we investigated if elevating the gluconate concentration could increase the naringenin production in ASA806. The 100 mM gluconate was used in the feeding medium, and the cultivation was performed for 102 h. As shown in Fig. 6, naringenin production was evident already after 6 h, reaching 52 mg/L within 31.5 h, corresponding to the productivity of 2.2 mg/L/h. The final titer of 66 mg/L was attained at the end of the cultivation. Thus, the fed-batch experiments also showed that most of naringenin was produced already during the first 31.5 hours, which is potentially related to the high availability of precursors in actively growing cells.

**Fig. 6.**
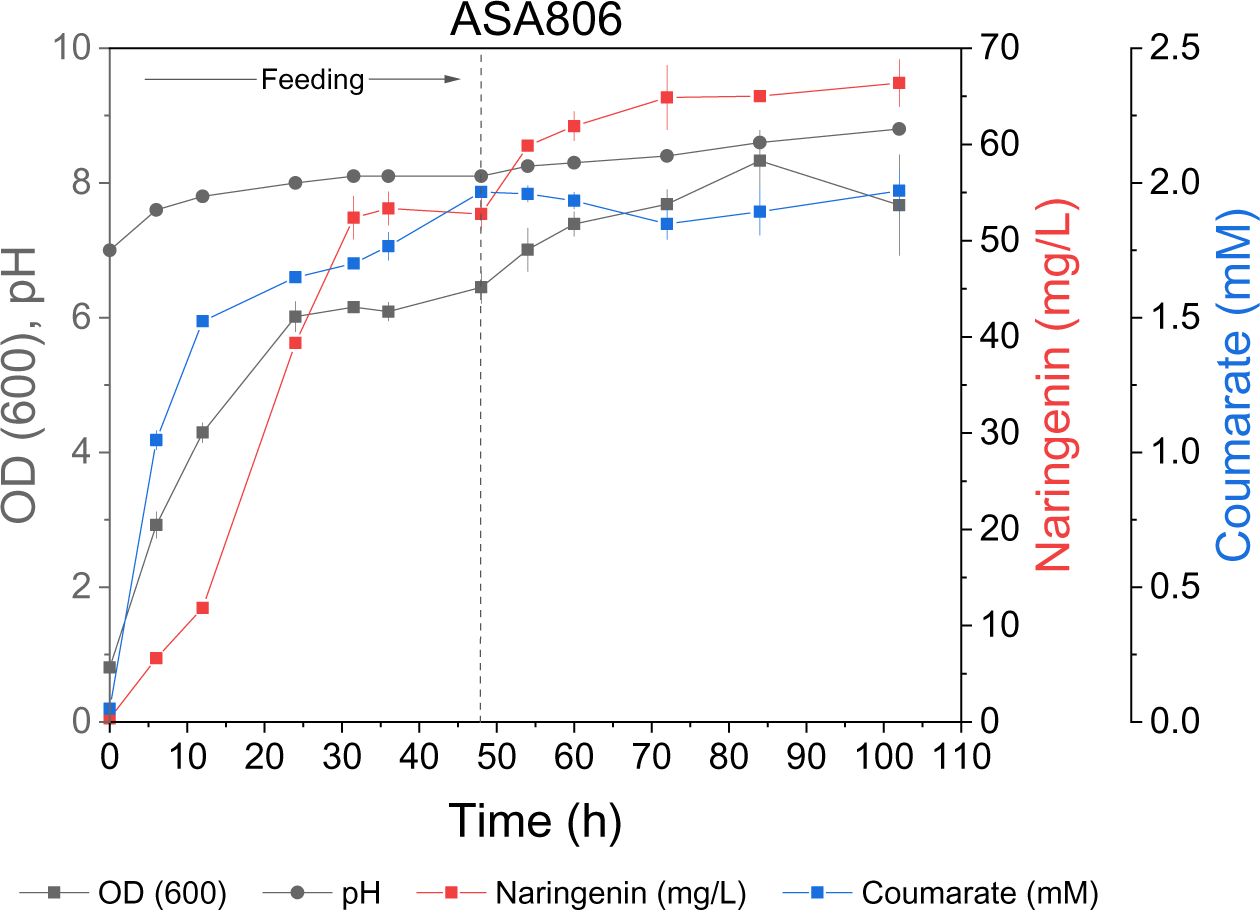
Fed-batch cultivation for naringenin production in higher concentration of gluconate. Cultures initially contained 50 μM of coumarate, 100 mM gluconate, and 0.2% casamino acids. Cultures were fed with a medium containing 100 mM gluconate, 0.2% casamino acids, and 3.2 mM coumarate for 48h (feeding rate 3.7mL/h). Error bars represent the mean ± s.d of two biological replicates.

In contrast to the study by Incha et al., (2020) who reported higher flaviolin with increased glucose supplementation, our engineered strain did not exhibit significantly higher naringenin production with increased supply of gluconate. This finding together with our results related to the experiments with engineered malonate pathway and cerulenin addition might indicate that the cellular malonyl-CoA content was not potentially limiting the naringenin synthesis in the studied conditions. It is also possible that the higher gluconate availability led to the direction of carbon towards the lipid production pathways of *A. baylyi* ADP1, which competes for acetyl-CoA through fatty acid synthesis (Luo et al., 2020; Arvay et al., 2021). This is further supported by the finding that the naringenin yield of ASA806 supplemented with 100 mM gluconate was 0.32 g_naringenin_/g_coumarate_, which was lower in comparison to a previous experiment with the same strain but with lower gluconate supplementation (0.54 g_naringenin_/g_coumarate_) (Table S3).

## 4. Conclusions

In this study, we engineered *A. baylyi* ADP1 for the production of naringenin from coumarate. We first eliminated the native catabolic pathway for coumarate utilization and then investigated several different combinations of CHS and CHI genes for the production. The highest naringenin titer, 66 mg/L, was obtained with the expression of CHS from *Hypericum androsaemum* and CHI from *Medicago sativa* in a fed-batch cultivation. The outcomes of our study demonstrate the potential of *A. baylyi* ADP1 as a host for flavonoid production. In particular, the high engineerability, the native aromatics catabolic pathways, and the natural ability to provide surplus of malonyl-CoA make *A. baylyi* ADP1 an attractive host for the production. While future research efforts are required to solve remaining issues related to insufficient catalytic efficiencies of CHS and CHI, and further increase of the malonyl-CoA pool for the naringenin synthesis pathway, the results achieved in this study mark a significant step towards the development of lignin-based production of value-added compounds in *A. baylyi* ADP1.

## Author contribution

KK, VS, and SS designed the study. KK carried out the microbiological and molecular work. EE and KK conducted HPLC analyses. All authors participated in writing the manuscript. All authors read and approved the final manuscript.

## CRediT authorship contribution statement

**Kesi Kurnia:** Conceptualization, Methodology, Validation, Visualization, Formal analysis, Investigation, Writing - original draft, Writing - review & editing. **Elena Efimova:** Methodology, Writing - original draft. **Ville Santala:** Conceptualization, Methodology, Writing - original draft, Writing - review & editing, Supervision. **Suvi Santala:** Conceptualization, Methodology, Writing - original draft, Writing - review & editing, Supervision, Funding acquisition.

## Acknowledgements

Funding: SS would like to thank the Novo Nordisk Foundation (grant NNF21OC0067758) and the Research Council of Finland (grant no. 334822 and 347204). VS would like to thank the Novo Nordisk Foundation (grant NNF21OC0079579).

## Appendix A. Supplementary data

